# Fluorescein Angiography Image-AI Based Early Diabetic Retinopathy Detection

**DOI:** 10.1101/2025.03.10.642348

**Authors:** Yiyan Peng, Huishi Toh, Peng Jiang

**Author notes:** To whom correspondence should be addressed: Peng Jiang.

## Abstract

Fluorescein angiography (FA) is widely regarded as the clinical gold standard for diagnosing diabetic retinopathy (DR), based on a range of observable symptoms, including microaneurysms, retinal hemorrhages, and vascular leakage. However, early detection of DR remains challenging, as these symptoms may not manifest simultaneously and are often too subtle to detect. Acellular capillary density is currently the only quantitative metric used to assess the risk and severity of DR, particularly in its early stages when these clinical symptoms are minimal or absent. Unfortunately, measuring acellular capillary density to detect pre-symptomatic diabetic retinopathy (DR) requires destructive techniques such as trypsin digest to remove non-vascular tissues, making it unsuitable for human patients, while current imaging techniques such as FA does not have resolution to quantify acellular capillary density, limiting the utility of non-invasive approaches for early detection of DR. In this study, we used Nile rat as a DR model to train an AI deep learning model based on FA images to detect retinas with acellular capillary density ≥18 counts per mm², which is considered as pre-symptomatic (before clinically observed DR symptoms) DR. Our model achieves an accuracy of 80.85%, with a sensitivity of 84.21%, specificity of 75.68%, and an area under the receiver operating characteristic (ROC) curve (AUC) of 0.86. We also showed that integrating the duration of diabetes as a pre-knowledge into a Bayesian framework with real time AI-image prediction can enhance the prediction power with AUC of ROC of 0.9. Using the first sequenced Nile rat genome and our RNA-seq data, we identified 43 gene markers that signified a shift from a normal state to a transitional phase preceding early diabetic retinopathy (DR). While PDGF-B/PDGFRβ signaling is typically associated with pericyte loss and increased acellular capillary density, which are hallmarks of early DR, none of the identified markers were related to this pathway. Instead, three genes (Bcl2a1, Birc5, and Il20rb) out of these 43 biomarkers are associated with inflammation or response to inflammation suggesting that inflammation may contribute to the shift in acellular capillary density from <16 (normal) to 17–18 counts (transitional phase preceding early DR) per mm² before pericyte loss occurs, providing new insights into the early pathogenesis of DR.

**Author Summary:** Diabetic retinopathy (DR) is a leading cause of vision loss worldwide. Early detection is critical to prevent progression, yet current diagnostic methods rely on symptoms that may appear too late. The density of acellular capillaries—tiny, non-functioning blood vessels in the retina—is a key early marker of DR, but measuring it requires destructive techniques that are not suitable for human patients. In this study, we used Nile rats, a rodent model that naturally develops diabetes and DR, to train an artificial intelligence (AI) model using fluorescein angiography (FA) images. Our AI system detects retinas with acellular capillary density at levels associated with pre-symptomatic DR with high accuracy. We further improved predictive power by integrating diabetes duration into a Bayesian framework. In addition, our transcriptomic study identified 43 markers associated with the transition from a normal state to a transitional phase preceding early diabetic retinopathy. Interestingly, none were linked to a well-known pathway involving pericyte loss, which has been thought to drive early DR. Instead, three markers (Bcl2a1, Birc5, and Il20rb) suggest inflammation may contribute to early changes before pericyte loss occurs. These findings offer a new perspective on the earliest stages of DR.

## Introduction

Diabetic retinopathy (DR) is a prevalent complication of diabetes that damages the blood vessels in the retina and can eventually lead to vision loss. In many developed nations, including the United States, it is the primary cause of new blindness cases among adults aged 20 to 74 [1]. Approximately one-third of individuals with diabetes, regardless of the subtype, are at risk of developing diabetic retinopathy[2].

Diabetic retinopathy [3–6] progresses through two main stages: non-proliferative DR and proliferative DR. Non-proliferative DR represents the early stage of the disease and is characterized by microvascular changes such as the formation of microaneurysms and the narrowing or closure of small blood vessels. At this stage, vision is typically unaffected. However, without timely intervention, non-proliferative DR often progresses to proliferative DR, a more severe stage marked by the growth of abnormal blood vessels in the retina and the formation of fibrous tissue. These fragile new blood vessels are highly prone to bleeding, leading to hemorrhages and significant vision loss. Early diagnosis of DR is therefore critical to prevent disease progression and preserve vision.

Fluorescein angiography (FA) is widely regarded as the clinical gold standard for diagnosing DR[7–9], based on a range of observable symptoms, including microaneurysms, retinal hemorrhages, and vascular leakage. However, early detection of DR remains challenging, as these symptoms may not manifest simultaneously and are often too subtle to detect. Currently, acellular capillary density is the only quantitative metric to assess the risk and severity of DR[10], particularly in its early stages when clinical symptoms are minimal or absent. Unfortunately, accurately measuring acellular capillary density requires destructive techniques such as trypsin digest to remove non-vascular tissues, making it unsuitable for human patients, while current imaging techniques such as FA does not have resolution to quantify acellular capillary density, limiting the utility of non-invasive approaches for early detection of DR.

The Nile rat (*Arvicanthis niloticus*), a diurnal rodent native to Northern Africa, offers a promising preclinical model for studying the progression of DR[10]. Unlike conventional laboratory rodents, Nile rats are highly susceptible to diet-induced diabetes when fed standard laboratory rodent chow, which is hypercaloric relative to their native diet [11]. This metabolic vulnerability mirrors the natural progression of type 2 diabetes in humans, making the Nile rat a valuable tool for studying diabetic complications. Notably, diabetic Nile rats develop DR with advanced retinal lesions resembling those observed in humans, including macular edema, capillary non-perfusion, and proliferative DR[10]. Prior studies [10, 12] have identified elevated acellular capillary density as a reliable early marker of DR, preceding vascular dysfunction in pericytes and endothelial cells. These findings underscore the significance of retinal vascular changes as a causal factor in the initiation and progression of DR.

In this study, we developed an AI FA-imaging based non-invasive diagnostic approach to detect early pre-symptomatic (before clinically observed symptoms) DR. We also showed that integrating the duration of diabetes as a pre-knowledge into a Bayesian framework with real time AI-image prediction can enhance the prediction power. Further transcriptomic analysis of Nile rat retina revealed 43 gene markers associated with normal to early DR transition phase. PDGF-B/PDGFRβ signaling is generally considered as a causal to pericyte loss and an increase in acellular capillaries—features typical of early diabetic retinopathy (DR). However, none of our identified markers were associated with this pathway. Instead, we detected inflammation or response to inflammation markers (Bcl2a1, Birc5, and Il20rb), which implies that inflammation might drive the rise in acellular capillary density from < 16 counts per mm² (normal) to 17–18 counts per mm² (transition phase before early DR) even before any pericyte loss occurs. This observation contributes to a debate over whether inflammation [13] or pericyte loss [14–16] is the primary initiator of the increased acellular capillary density that heightens DR risk. Our data suggest that inflammation may indeed play earlier role in the initial progression of DR potentially before pericyte loss, offering new insights into the early pathogenesis of DR.

## Results and Discussions

### Diabetes Duration as a Robust Predictor of Elevated Acellular Capillary Density

In our previous study [12], we analyzed a small sample size of Nile rats (N=32) to identify risk factors associated with increased acellular capillary density. We found elevated random blood glucose and a high-caloric, low-fiber diet each contributed to acellular capillary density increase. Sex had only a marginal effect, mainly in individuals with both high glucose and a high-caloric diet [12]. In this study, we expanded our sample size to 124 Nile rats (N=124) to more reliably identify key risk factors associated with elevated acellular capillary density. We employed a random forest regression model to assess feature importance and rank each factor based on its contribution. The model was trained using diabetes duration, mean random blood glucose (RBG), diabetes status (diabetic vs. non-diabetic), diet (high- vs. low-fiber), age, and sex as input variables, with acellular capillary density as the dependent variable. Model performance was evaluated using mean squared error (MSE), which measures the average difference between predicted and actual values. Feature importance was assessed using mean decrease in accuracy (MDA), which quantifies the increase in prediction error when a given variable is randomly permuted. A higher MDA indicates a stronger contribution of that feature to the model’s predictive power, with larger values reflecting greater importance in determining acellular capillary density. As shown in **Figure 1A**, diabetes duration, mean RBG, diabetes status, diet, and age are all associated with increased acellular capillary density, while sex has the least impact. Among these factors, diabetes duration emerged as the strongest and most consistent predictor of acellular capillary density. The correlations between diabetes duration and acellular capillary density are shown in **Figure 1B**. No significant sex differences were observed when diabetes duration was less than 20 weeks. However, after 20 weeks, males exhibited a slightly greater increase in acellular capillary density compared to females, suggesting that sex may act as a modest conditional modifier dependent on diabetes duration. Nevertheless, diabetes duration remains the most robust and reliable risk factor for increased acellular capillary density. This is consistent with findings in humans, where the duration of diabetes is a well-established risk factor for diabetic retinopathy [17].

**Figure 1.**
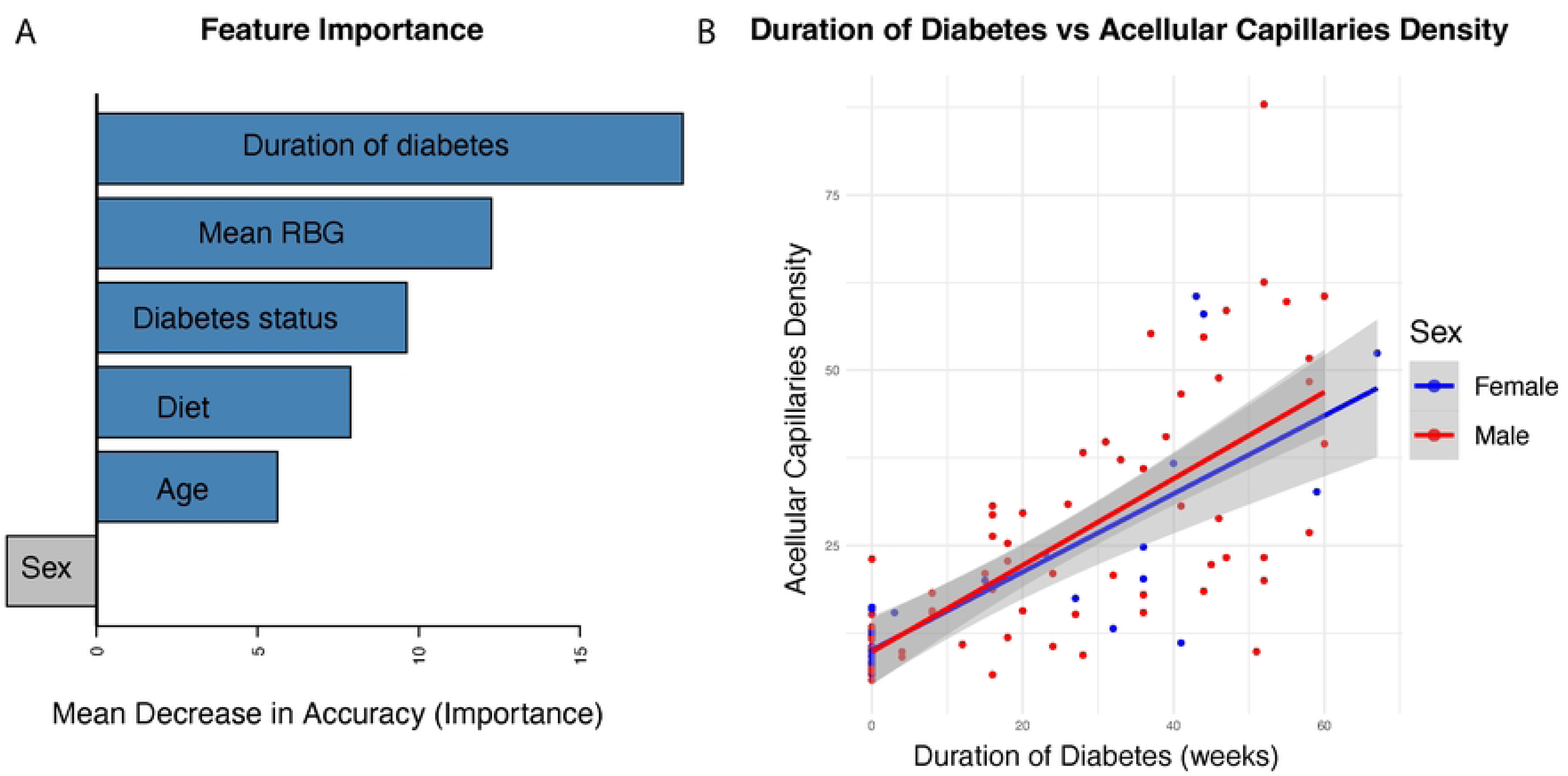
Factors influencing acellular capillary density. (A) Feature importance ranking based on the random forest regression model. (B) Association between diabetes duration and acellular capillary density.

### Deep Learning Model for Early Detection of Diabetic Retinopathy

Fluorescein angiography (FA) is widely recognized as the clinical gold standard for diagnosing diabetic retinopathy (DR) by identifying hallmark symptoms such as microaneurysms, retinal hemorrhages, and vascular leakage. However, for early pre-symptomatic DR, FA has limitations in resolving subtle changes in retinal vascular morphology. We hypothesized that a deep learning AI model could detect these early, clinically pre-symptomatic vascular changes by identifying complex patterns in FA images that may not be readily visible through conventional analysis. To test this hypothesis, we developed an AI-based classification model using Densely Connected Convolutional Networks (DenseNet-169) to predict acellular capillary density ≥18 counts per mm² (high DR risk) versus <18 counts per mm² (low DR risk) from FA images. We defined a cutoff of 18 counts per mm² based on prior findings indicating that densities below 16 counts per mm² are unlikely to be associated with retinopathy[10, 12]. Our analysis of 124 Nile rats revealed a significant increase in both random blood glucose levels and diabetes duration when comparing animals with acellular capillary densities of 10–16 counts per mm² versus 20–22 counts per mm². Notably, median random blood glucose increased from 123 mg/dL to 209.4 mg/dL (Wilcoxon test, P = 0.0221), and median diabetes duration increased from 0 to 28 weeks (Wilcoxon test, P = 0.00801), suggesting a sharp rise in retinopathy risk as capillary density surpasses 16 counts per mm² (**Supplementary Figure S1**). Based on this trend, we used 18 counts per mm² as the threshold for defining early diabetic retinopathy, even in the absence of typical symptoms such as microaneurysms, retinal hemorrhages, and vascular leakage. This threshold marks the transition point where the likelihood of DR progression significantly increases. Using this cutoff, our AI model classifies the FA image as high DR risk (≥18 counts per mm²) or low DR risk (<18 counts per mm²).

DenseNet-169 is a deep convolutional neural network known for its densely connected layers, where each layer receives feature maps from all preceding layers, enhancing feature propagation and reducing the number of parameters. In this study, we applied transfer learning by leveraging the pre-trained weights of DenseNet-169 which were originally trained on large-scale image datasets, fine-tuning the last convolutional block and the fully connected classification layer to adapt the model for our classification task, while keeping the earlier feature extraction layers unchanged (**Figure 2**). This approach allowed the model to retain general image feature representations while learning to classify acellular capillary density as exceeding (high DR risk) or falling below (low DR risk) 18 counts per mm². As shown in **Figure 3**, the deep learning model can achieve an overall accuracy of 80.85%, with a sensitivity of 84.21%, specificity of 75.68%, and an area under the receiver operating characteristic (ROC) curve (AUC) of 0.86 via a 5-fold cross-validation. Among the total images analyzed, the model correctly identified 48 true positives and 28 true negatives, with 18 incorrect predictions. The robust polynomial fitting curve (Figure 3), generated from correctly predicted cases, suggests that the 15 to 23 counts per mm² range may represent a critical transition window for early DR classification. Within this range, AI-predicted probability aligns strongly with observed trypsin digest–based acellular capillary density counts. This finding suggests that fluctuations in acellular capillary density within this window could be biologically significant for early DR progression. Beyond this range—above 23 or below 15 counts per mm²—the AI-predicted probability does not show a strong correlation with observed density values. Beyond the 15 to 23 counts per mm² range, AI-predicted probability does not increase or decrease proportionally with higher or lower acellular capillary densities. Specifically, when acellular capillary density exceeds 23 counts per mm² or falls below 15 counts per mm², the AI-predicted probability reaches a plateau, suggesting a potential saturation effect or insufficient sensitivity in these extreme ranges.

**Figure 2.**
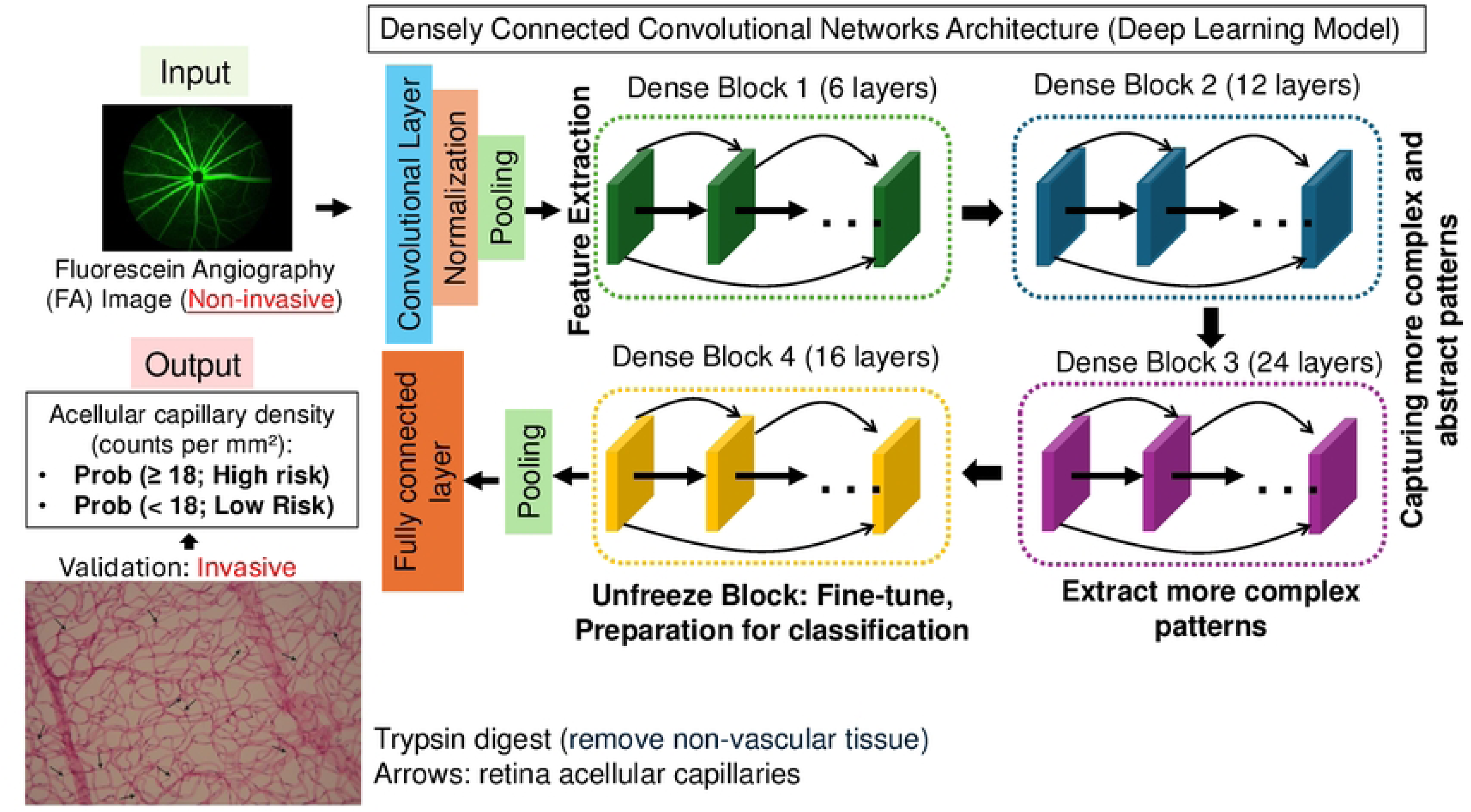
Deep learning model for early retinopathy prediction using fluorescein angiography (FA). A DenseNet-169-based AI model was trained on FA images to predict retinopathy risk based on acellular capillary density (≥18 per mm²: high risk; <18 per mm²: low risk). The ground-truth (acellular capillary density) was based on trypsin digest (invasive) to count acellular capillaries (arrows).

**Figure 3.**
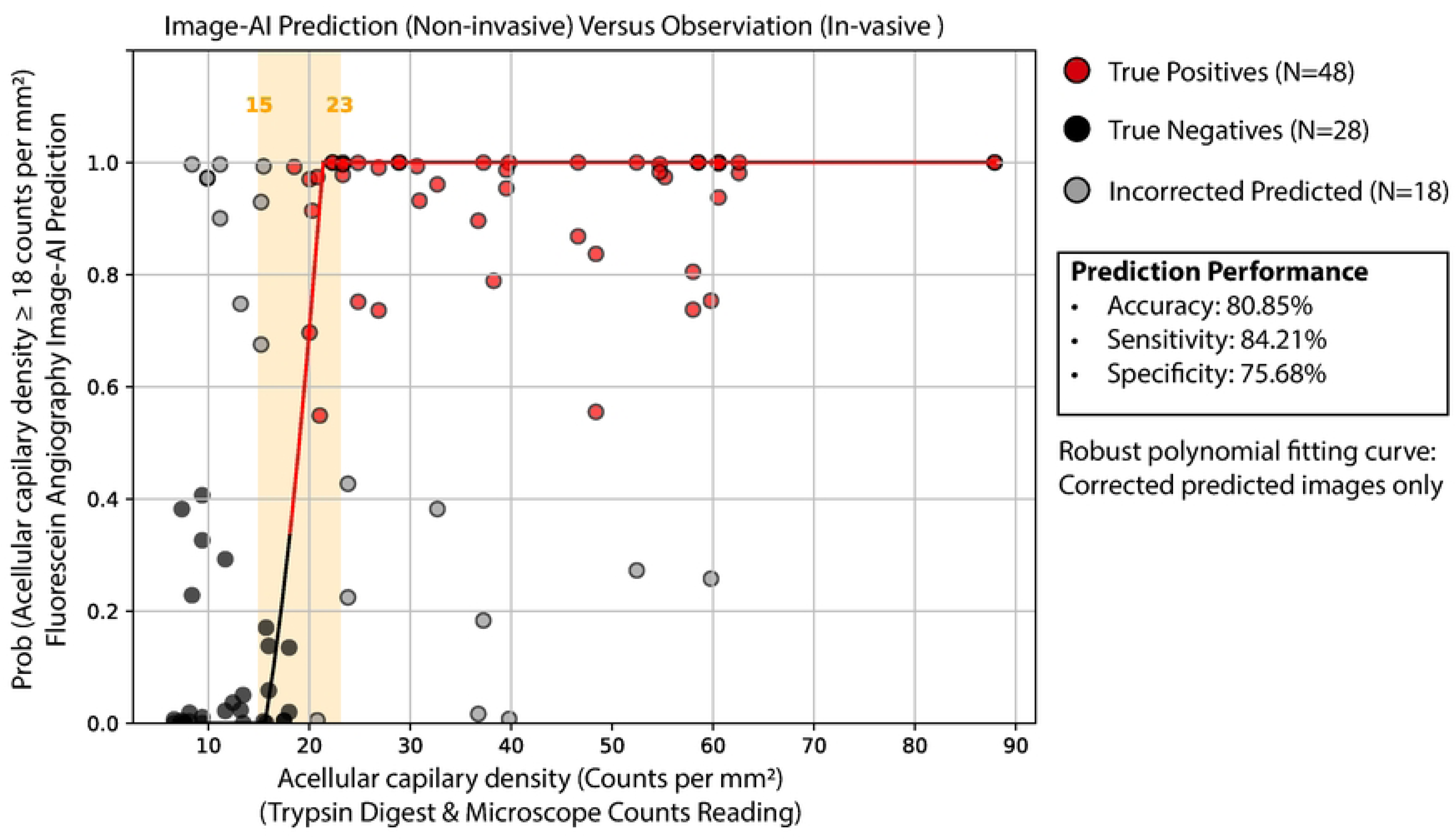
AI-predicted vs. ground truth acellular capillary density. Comparison of FA-based AI predictions with microscopic counts from trypsin digest. The model achieved 80.85% accuracy, 84.21% sensitivity, and 75.68% specificity.

These findings reinforce the hypothesis that the 15 to 23 counts per mm² range represents a critical transition window in early DR progression. Within this range, acellular capillary density fluctuations may reflect key pathological changes that are detectable by AI classification models but not easily captured in a continuous, quantitative manner. This underscores the potential importance of this range for early DR risk assessment and suggests that further investigation into its biological significance could enhance early diagnostic strategies.

### Integration the Duration of Diabetes with AI-based Real-time Image Prediction Enhanced Early Diabetic Retinopathy Detection

A key advantage of an AI-based FA image retinopathy detection model is its ability to provide real-time predictions. While the duration of diabetes is a well-established and significant risk factor for diabetic retinopathy, not all diabetic patients develop retinopathy, and not all individuals with DR have a long history of diabetes. In humans, one-third of individuals with diabetes will develop DR [2]. More importantly, the timing of DR onset varies among individuals, and the recorded duration of diabetes is often inaccurate in real clinical settings. Intuitively, if a patient with a long history of diabetes receives a high DR risk prediction from the AI model, this prediction is more likely to be accurate compared to a non-diabetic individual receiving the same DR risk (where the AI prediction might be incorrect). Based on this reasoning, we hypothesize that incorporating the duration of diabetes as a prior and integrating this prior knowledge into a Bayesian framework to calculate the posterior probability of AI-based DR predictions could enhance prediction accuracy. Specifically, we used logistic regression to model the relationship between diabetes duration and the probability of acellular capillary density ≥18 counts per mm². This probability was then incorporated into a Bayesian formula (see Methods) to compute the posterior probability of real-time AI image-based DR prediction. As shown in **Figure 4A**, incorporating the duration of diabetes as a prior with the image AI model resulted in improved classification performance compared to the image AI model alone. The AUC of the ROC curves for the AI-based model alone and Bayesian model incorporating diabetes duration achieves are 0.86 and 0.90, respectively. This finding suggests that combining prior knowledge (duration of diabetes) with real-time AI-based image predictions can enhances the reliability and robustness of early retinopathy detection.

**Figure 4.**
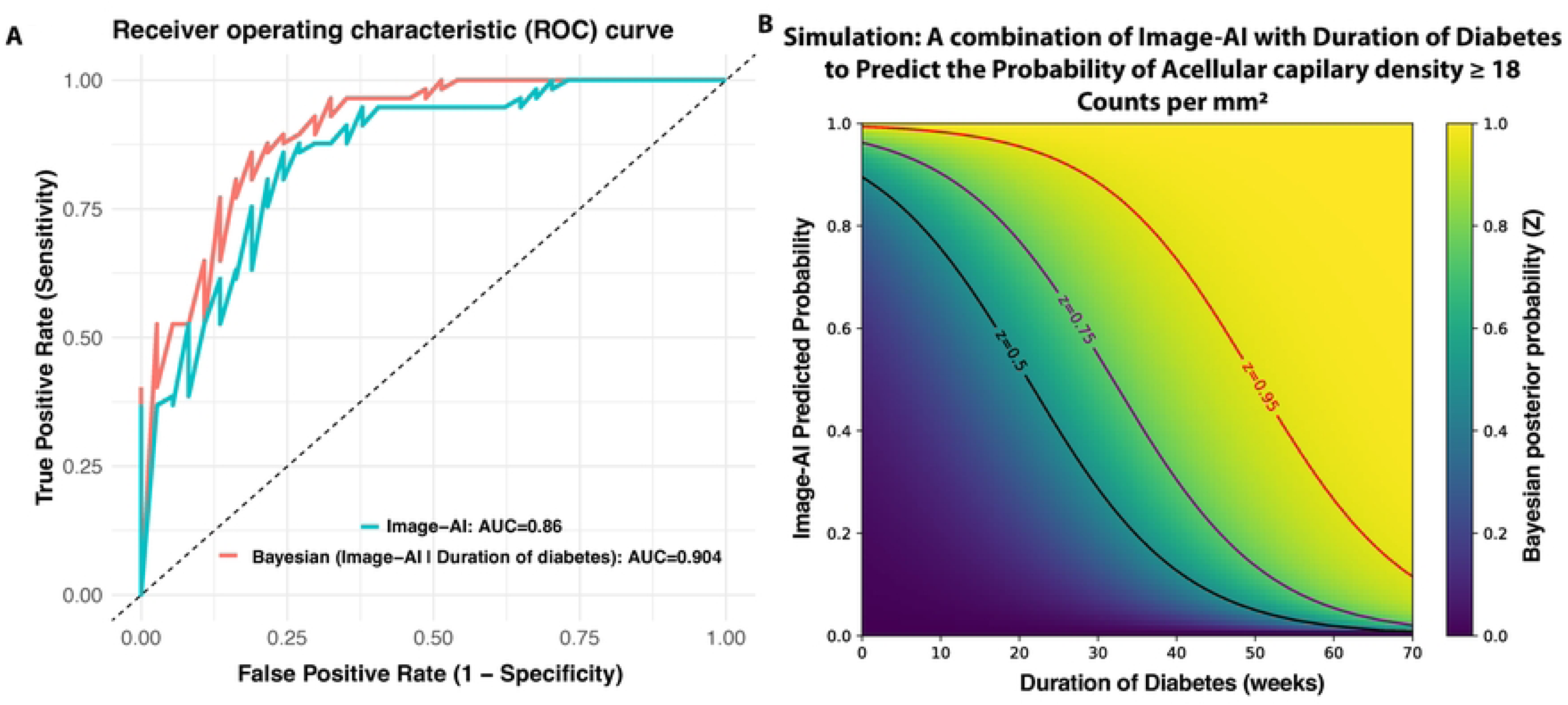
Bayesian framework integrating diabetes duration with AI predictions. (A) Improved prediction accuracy using a Bayesian approach. (B) Simulation of how diabetes duration combined with AI predictions influences posterior probability of acellular capillary density ≥18 counts per mm².

We further conducted a simulation study to evaluate all possible combinations of diabetes duration (prior knowledge) and real-time AI-based image predictions in generating posterior probabilities for retinopathy. As shown in **Figure 4B**, when the duration of diabetes is short, the AI model must assign a very high probability to achieve a high Bayesian posterior probability. In contrast, as the duration of diabetes increases, the posterior probability remains high even when the AI-predicted probability is moderate. This suggests that for individuals with a shorter duration of diabetes, the AI model must produce a very high probability to predict retinopathy accurately. However, for those with a longer duration of diabetes, even moderate AI-predicted probabilities are sufficient to yield high posterior probabilities. This relationship highlights the importance of incorporating prior knowledge of diabetes duration to refine AI-based predictions. By integrating this Bayesian approach, clinicians can make more informed decisions, particularly in borderline cases where AI predictions alone may be uncertain. In practice, this framework could help prioritize follow-up screenings, guide early intervention strategies, and improve risk stratification for diabetic retinopathy, ultimately enhancing patient outcomes.

### Detection of Biomarkers Associated with Shifting From a Normal State to a Transitional Phase preceding Early Diabetic Retinopathy

Although the Nile rat is a valuable preclinical model for diabetic retinopathy, closely mimicking human disease with progressive retinal vascular lesions such as increased acellular capillaries, pericyte loss, microglial infiltration, vascular leakage, capillary non-perfusion, and neovascularization[10], RNA-seq-based transcriptomic studies remain highly limited. This is because standard RNA-seq workflows require mapping reads to a fully sequenced and annotated genome or transcriptome. However, until recently, no complete genome was available for the Nile rat. To solve this problem, we developed CRSP (comparative RNA-seq pipeline) [18], a tool that integrates multiple comparative cross-species computational strategies, including novel transcript assembly and mapping to annotated mouse transcripts, to impute gene expression levels in the Nile rat. While this comparative species computational approach is valuable, it also introduces potential data noise and technical bias due to differences in genome architectures, sequence divergence, and incomplete cross-species transcript annotation, which may lead to inaccuracies in gene expression quantification. Recently, we released the first fully sequenced and annotated Nile rat genome [19], which could serve as a direct mapping reference. However, like any newly sequenced genome, it still contains unsequenced regions and potential annotation errors. To determine whether our comparative cross-species pipeline or the newly sequenced genome provides more reliable gene expression estimates, in this study we conducted the first benchmarking study to evaluate these two strategies based on our prior RNA-seq of 28 Nile rat retinal vasculature samples [12].

We first calculated the Spearman’s rank correlation coefficient (Rho) between transcripts per million (TPMs; gene expression values) estimated by CRSP (comparative RNA-seq pipeline), which was used in our prior study, versus TPMs obtained from mapping reads to the newly sequenced and annotated Nile rat genome. The correlation was moderate (Rho = 0.37), suggesting a substantial discrepancy between these two computational approaches. Given that RNA-seq read count distributions are often modeled using a negative binomial (NB) distribution [20–22], we next examined whether gene read counts estimated by CRSP and genome mapping both followed the expected NB distribution. In the NB framework, the overdispersion ratio is used to quantify the extent to which variance exceeds the mean. A higher overdispersion ratio indicates greater variability in gene expression, often due to technical noise. In contrast, a lower overdispersion ratio suggests more stable gene expression estimates, with variance closer to the mean, making the data less affected by unwanted variability. To assess this, we calculated the overdispersion ratio for both approaches. We found that transcript read counts estimated by mapping to the newly sequenced Nile rat genome had a statistically significantly lower overdispersion ratio compared to those estimated using CRSP (**Figure 5B**). This suggests that RNA-seq expression estimates obtained from direct genome mapping exhibit greater reliability than those generated using the CRSP strategy, which relies on cross-species computational imputation and may introduce additional noise due to sequence divergence and incomplete transcript annotation.

**Figure 5.**
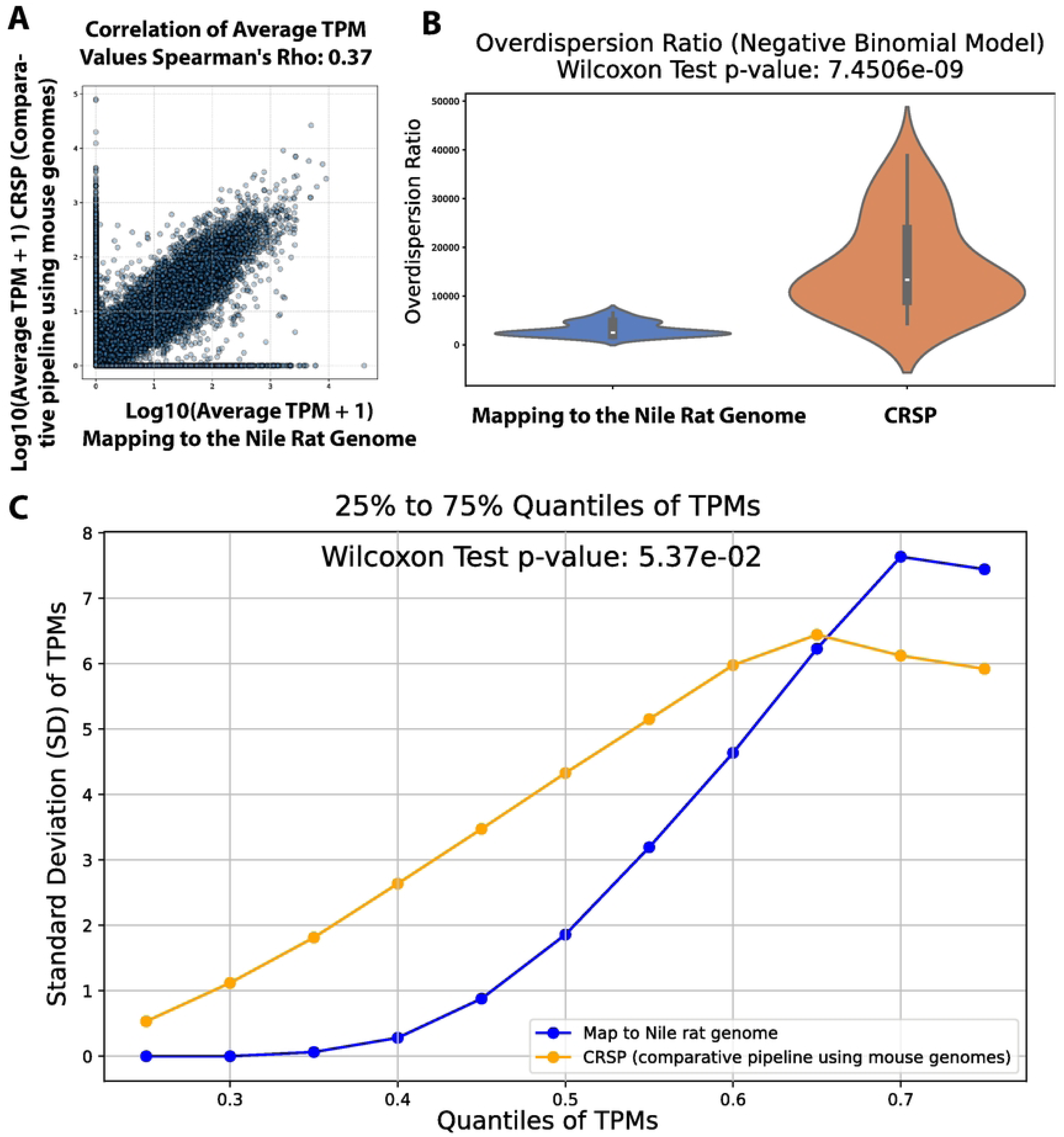
Comparison of gene expression estimates from RNA-seq reads mapping to Nile rat genome vs. CRSP tool (comparative-species pipeline does not rely on a genome). (A) Correlation of gene expression values between methods. (B) Mapping to the Nile rat genome results in significantly lower over-dispersion than CRSP. (C) Comparing the standard deviations of gene expression within the same quantile between these two methods.

To investigate this further, we reasoned that by sorting gene expression values (TPMs) from low to high within each RNA-seq sample and grouping genes across samples based on the same quantile (from 25% to 75% with 5% increments), we could minimize biological variation. This quantile-based approach ensures that variations observed across samples within the same quantile are primarily driven by technical factors rather than biological differences, as genes are ranked solely by expression level. As shown in **Figure 5C**, when we compared the standard deviations of TPMs within each quantile, gene expression values derived from mapping to the Nile rat genome exhibited significantly lower standard deviations than those estimated using CRSP. This suggests that using the newly sequenced Nile rat genome as a mapping reference yields more consistent gene expression estimates than the CRSP strategy we previously employed.

Therefore, we utilized gene expression data derived from mapping to the newly sequenced Nile rat genome to identify biomarkers associated with the pre-symptomatic transition phase of diabetic retinopathy. In non-diabetic Nile rats, acellular capillary density typically ranges from 5.8 to 16.2 counts per mm² [10], with densities below 16 counts per mm² generally considered normal. However, when acellular capillary density reaches ≥18 counts per mm², it is classified as high-risk for retinopathy due to a marked increase in both random blood glucose levels and diabetes duration. This suggests that the 16 to 18 counts per mm² range represents a critical transition phase in the progression of diabetic retinopathy. To identify potential gene biomarkers associated with this transition, we conducted a differentially expressed genes (DEG) analysis, comparing normal (<16 counts per mm²) and transition (17–18 counts per mm²) samples. **Figure 6A** presents the retinal vascular RNA-seq samples used for biomarker identification. We identified 43 DEGs (all of them are up-regulated) between normal and transition samples (Figure 6B), using a threshold of >2-fold changes in normalized read counts with a false discovery rate (FDR) <5%. Notably, among these 43 differentially expressed genes (DEGs), three genes (Bcl2a1, Birc5, and Il20rb) are associated with inflammatory response. Several studies showed that Bcl2a, Birc5 and Il20rb were associated with chronic inflammation and autoimmune diseases[23–26], suggesting that inflammation could be one of the factors to drive the early onset of DR.

**Figure 6.**
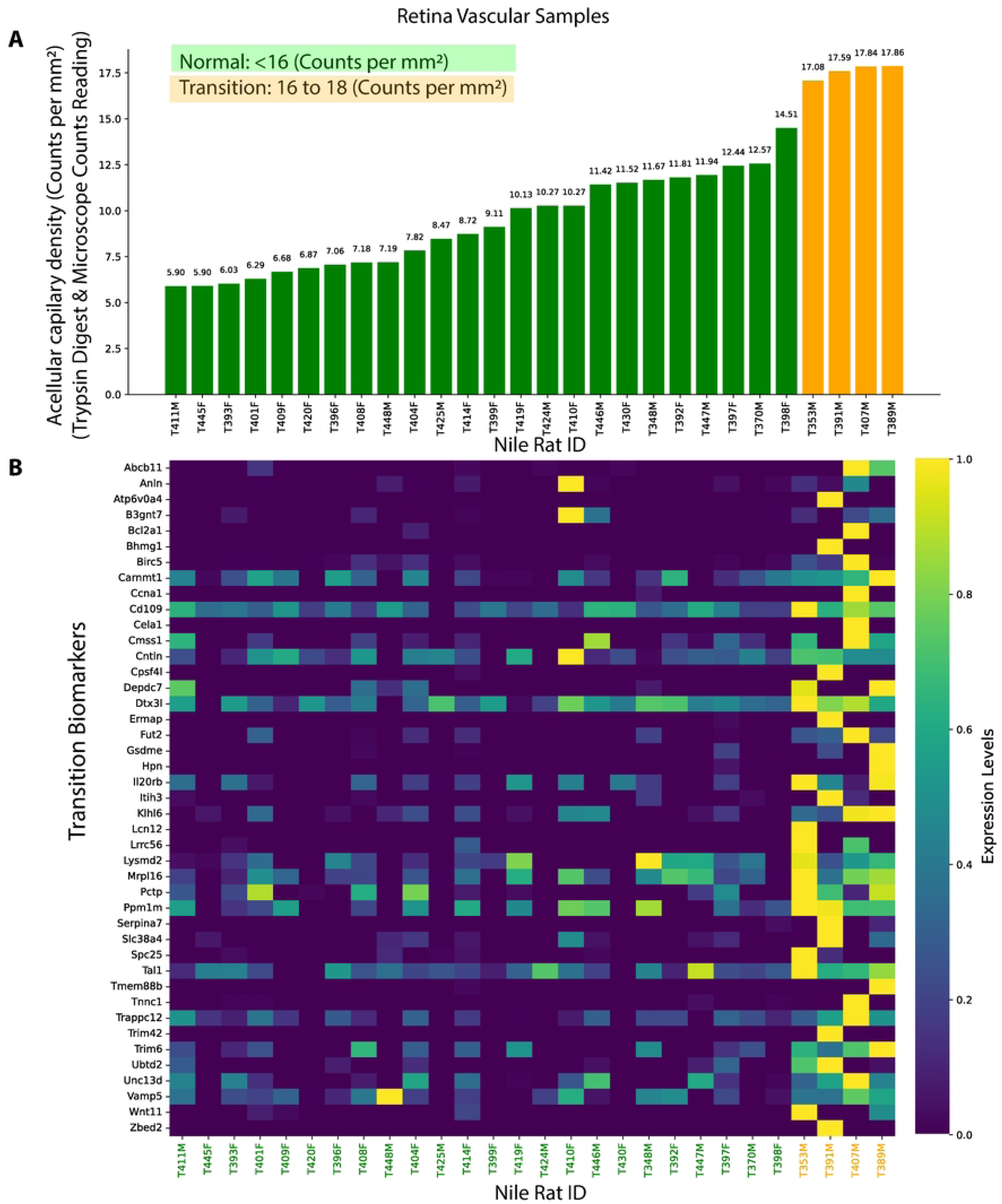
Retinal gene markers associated with early diabetic retinopathy (DR). (A) Nile rat RNA-seq samples used for gene marker identification, with green indicating normal and orange representing transition-phase samples before early DR. (B) Identified gene markers distinguishing normal and transition-phase samples.

Increased acellular capillary density is considered as one of the measurable signs of the onset diabetic retinopathy [10, 27]. The primary cause of acellular capillaries is the loss of pericytes from the capillary walls, resulting in the formation of non-functional “empty” basement membrane tubes, also known as ghost vessels. Pericytes play a crucial role in maintaining the blood-retina barrier, regulating endothelial proliferation, and stabilizing retinal capillaries. Among the 43 biomarkers analyzed during the transition from normal to early diabetic retinopathy (DR), we did not identify any genes associated with PDGF-B/PDGFRβ signaling. However, the presence of inflammation markers (Bcl2a1, Birc5, and Il20rb) suggests that the shift in acellular capillary density—from less than 16 counts per mm² (typically considered normal) to 17–18 counts per mm² (indicative of transition)—could be driven by inflammation as one of the contributing factors potentially even preceding pericyte loss. In fact, several studies found an association between diabetic retinopathy and chronic inflammation, autoimmune diseases, such as rheumatoid arthritis [13], suggesting inflammatory conditions may exacerbate retinal microvascular dysfunction and contribute to the early pathogenesis of diabetic retinopathy.

## Conclusions

Early detection of diabetic retinopathy (DR) remains challenging due to the absence of clear clinical symptoms in its initial stages. Our study addresses this gap by leveraging Nile rats as a DR model to train an AI deep learning system using fluorescein angiography (FA) images, achieving high accuracy in detecting retinas with acellular capillary density ≥18 counts per mm²—a threshold indicative of pre-symptomatic DR. Integrating diabetes duration into a Bayesian framework further improved predictive power. Additionally, transcriptomic analysis identified 43 gene markers associated with the transition to pre-symptomatic DR, with inflammation-related genes (Bcl2a1, Birc5, and Il20rb) emerging as potential drivers of early vascular changes. These findings suggest that inflammation, may play a key role in the initial the early progression of DR even before pericyte loss [14–16], providing new insights into its early pathogenesis.

## Methods

### Animal protocol and ethical Statement

All animal experiments were approved by the University of California Santa Barbara, Institutional Animal Care and Use Committee, and conducted in accordance with the NIH Guide for the Care and Use of Laboratory Animals; study protocol 893. UCSB founder Nile rats were derived from the Brandeis University colony of the KC Hayes Laboratory. Nile rats in UCSB are housed at 21– 26°C in a conventional facility with individually ventilated cages and are provided autoclaved Sanichips as bedding material. Nile rats are either fed a high-fiber diet (Lab Diet 5L3M; Newco Speciality, Rancho Cucamonga, CA, USA) or a high-caloric diet (Formulab Diet 5008; Newco Speciality, Rancho Cucamonga, CA, USA) [28]. The percentage of crude fiber is 23% for the high-fiber diet and only 4% for the high-caloric diet. The ratio for the percentage of calories provided by carbohydrate, fat, and protein were 67:10:23 for the high-fiber diet and 56:17:27 for the high-caloric diet. Random blood glucose (RBG) levels are measured every four weeks starting at weaning age (4 weeks old). To reduce adverse events from diabetic complications, Nile rats with RBG>500 mg/dL were euthanized.

### Retinal trypsin digest and acellular capillary count reading

From each animal, we did a trypsin digest on retina to count acellular capillary density. The retinal vasculature was loosened from other retinal cells using osmotic lysis, where the retina was placed in RNase-free water for 1h at 4°C on a shaker. Next, water was aspirated, and the remaining retinal tissue was incubated in a 10% DNase solution at 37°C for 15 minutes. Then, the retina was transferred to a petri-dish with RNase-free water. Working under a dissecting microscope, we used a P200 pipet to squirt water on the retina until the inner limiting membrane was detached and neuronal cells were washed away from the retinal vasculature. For the trypsin digest, we used our previously published method [10]. Micrographs for quantification were taken with a Canon Rebel XSi digital camera (Canon, Tokyo, Japan) attached to an Olympus CKX41 microscope via an LM scope C-mount. For acellular capillary counts, 12 micrographs were taken from 3 randomly selected areas in each of the 4 retinal quadrants and analyzed at a final magnification of 200×. The total area used for quantification corresponded to approximately 8.5% of the whole retinal area. The counts were quantified using FIJI computer software by two masked graders.

### Defining pre-symptomatic retinopathy

Acellular capillary density below 10 counts per mm² has been considered very unlikely to be associated with retinopathy [12] and is considered normal. In prior study [10], the acellular capillary counts per mm² in Nile rats without diabetes reached as high as 16.2, suggesting that densities ranging from 10 to 16 counts per mm² are still less likely to be linked with retinopathy. In this study, based on 124 Nile rats, we observed a significant shift in both random blood glucose levels and diabetes duration when comparing the group with 10 to 16 counts per mm² to the group with 20 to 22 counts per mm² (**Supplementary Figure S1**). Specifically, the median random blood glucose increased from 123 mg/dL in the lower density group to 209.4 mg/dL in the higher density group (Wilcoxon test, P-value = 0.0221), while the median duration of diabetes increased from 0 weeks to 28 weeks (Wilcoxon test, P-value = 0.00801). This indicates a sharp increase in retinopathy risk when the acellular capillary density increases from 16 to 20 counts per mm². Based on this pattern, we define the midpoint (18 counts per mm²) as the cutoff for pre-symptomatic retinopathy, marking the transition point where the likelihood of developing retinopathy begins to rise substantially.

### Fluorescein angiography (FA) image data collection

After brief anesthesia with isoflurane, the Nile rats were injected (IM) with Midazolam (3 mg/kg; Akorn Inc., Lake Forest, IL, USA). Standard procedures were followed using the Phoenix MicronIV (Phoenix Research Laboratories, Pleasanton, CA) to perform fundus fluorescein angiography (FFA). We used 1% tropicamide ophthalmic solution, 2.5% phenylephrine hydrochloride ophthalmic solution (Paragon BioTeck, Inc. Portland, Oregon, USA), GONAK Hypromellose Ophthalmic Demulcent Solution, and AK-Fluor 10% fluorescein (intraperitoneal injection, 1 μl/g body weight). All ophthalmic solutions and fluorescein were from Akorn Inc. (Lake Forest, IL, USA), unless otherwise indicated.

### Fluorescein angiography (FA) image based AI model for early diabetic retinopathy detection

We developed an AI model to detect early diabetic retinopathy using fluorescein angiography (FA) images. The dataset consisted of 94 images (N=94) from 51 Nile rats, with 57 classified as diabetic retinopathy (acellular capillary density ≥18 counts per mm²) and 37 as control (acellular capillary density <18 counts per mm²). FA images underwent preprocessing, including resizing to 224 pixels and center cropping, to ensure consistent input dimensions. All images were normalized using mean values of [0.485, 0.456, 0.406] and standard deviations of [0.229, 0.224, 0.225]. The dataset was split into training and validation sets using five-fold cross-validation (K = 5). Each fold used 80% of the data for training and 20% for validation, ensuring that all samples were included in both training and validation across different iterations. Data augmentation was only applied to the training set, incorporating random rotation (10°) and horizontal flipping to enhance model generalization. The validation set was processed without augmentation.

The model was based on DenseNet-169, which consists of four dense blocks and a classifier. Transfer learning was applied by freezing all layers except for the last dense block (DenseBlock4) and the classifier. Fine-tuning was performed to optimize feature extraction for retinopathy classification. The existing classifier was modified to include a dropout layer (rate = 0.5) for regularization and a fully connected layer with two output neurons for binary classification. The model was trained using the Adam optimizer with cross-entropy loss to measure the discrepancy between predicted and true labels. Hyperparameter optimization was conducted by testing different combinations of epoch numbers, initial learning rates, regularization strengths, batch sizes, and step sizes. The model that achieved the best performance, with an overall accuracy of 80.85%, used the following hyperparameters: epoch number = 30, batch size = 10, initial learning rate = 0.01, and weight decay (regularization strength) = 1e-4. A step learning rate scheduler reduced the learning rate by a factor of 0.1 every five epochs. During the validation phase, probabilities for each class were computed using the Softmax function (which, for binary classification with two output neurons, is conceptually equivalent to logistic regression) and stored along with the predictions for further analysis. Accuracy for each fold was calculated as the percentage of correct predictions, and the overall accuracy was obtained by averaging the validation accuracies across the five folds to assess the model’s overall performance. The model was implemented using Python (version 3.12.3) and PyTorch (version 2.5.0).

### A Bayesian framework to integrate the duration of diabetes with real time FA-image AI model to predict DR

We developed a Bayesian framework that integrates duration of diabetes with a real-time fluorescein angiography (FA) image-based AI model to improve diabetic retinopathy (DR) prediction. First, we used logistic regression to model the relationship between the duration of diabetes (D) and acellular capillary density, estimating the probability of DR (acellular capillary density greater than 18 counts per mm²) as follows:

The prior probability is given by:

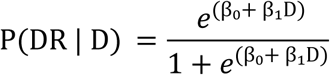

where:

- P(DR∣D) is the probability of diabetic retinopathy (DR), defined as acellular capillary density greater than 18 counts per mm²) given the duration of diabetes D.
- β₀ and β₁ are regression coefficients.
- D represents the duration of diabetes.

The maximum likelihood estimation (MLE) is used to find the values of β₀ and β₁ of the logistic regression model to maximize the probability of observing the given data. We convert DR into binary outcomes (0 = no DR, 1 = DR). The likelihood function is:

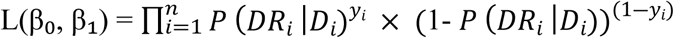

The likelihood function is further log10 transformed as:

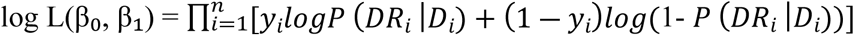

where:

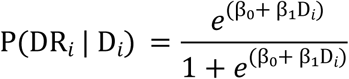

Given a particular Nile rat, the P(DR∣D) which is considered a prior probability will be estimated. We then used a Bayes’ theorem to combine the image-AI predicted DR probability, denoted as P(DR | AI) with the prior probability P(DR∣D)

Then, the AI-image predicted DR probability, denoted as P(DR | AI), will be integrated with the prior probability P(DR∣D) via a Bayes’ theorem to calculate the posterior probability of DR:

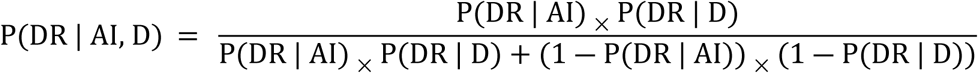

where P(DR | AI, D) represents the posterior probability of DR considering both the duration of diabetes and AI-based imaging predicted probability.

### RNA-seq data analysis

The raw RNA-seq FASTQ files from retina vascular biopsy samples were from our previous study [12] and are available under the GEO accession number GSE220672. We re-analyzed the raw RNA-seq data via mapping to the newly sequenced and annotated Nile rat genome [19]. Specifically, RNA-seq reads were mapped to the Nile rat annotated protein-coding genes using Bowtie [29], allowing up to 2-mismatches. The gene expected read counts and Transcripts Per Million (TPM) were estimated by RSEM [30]. The gene expected counts were further normalized by median-of-ratio method via EBSeq [31] R package. The re-analyzed RNA-seq data via mapping to the Nile rat annotated protein-coding genes was also compared with gene expression values (TPMs and mapping counts) estimated using our previous CRSP (Comparative RNA-seq Pipeline) tool, which does not rely on a sequenced genome. Biomarker detection was conducted based on gene expression values derived from mapping to annotated protein-coding genes in the Nile rat. The EBSeq package [31] was used to assess the probability of gene expression being differentially expressed (biomarkers) between acellular capillary density range from <16 (normal) to 17–18 counts (transitional phase preceding early DR) per mm². We required that differentially expressed genes should have false discovery rate (FDR)<5% via and >2 fold-change of median-by-ratio normalized read counts.

## Availability of the computational codes

The python codes to develop the Image-AI model is available in **Supplementary File 2**.

## Acknowledgments

We thank the technical support for the AI component from Dr. Le Yang from Kelvin Innovations LLC, Madison, WI, 53705, USA. We also thank the support from the Center for Gene Regulation in Health and Disease (GRHD) at Cleveland State University and Prof. Anton Komar.

## Funding

This study was supported by the Garland Initiative for Vision and funded by the William K. Bowes Jr. Foundation.

## List of Supplementary Data

**Supplementary Figure S1:** Comparison of samples with acellular capillary density of 10–16 counts per mm² and 20–22 counts per mm². Significant differences were observed in mean random blood glucose (A) and diabetes duration (B) between these groups, supporting our classification of the 16–20 counts per mm² range as a critical transition window from normal to early diabetic retinopathy. This rationale justifies selecting the median value (18 counts per mm²) within this range to define early diabetic retinopathy.

**Supplementary File 2**: Python codes for the Image-AI model

## Notes

### Competing Interest Statement

The authors have declared no competing interest.

